# On Pre-trained Language Models for Antibody

**DOI:** 10.1101/2023.01.29.525793

**Authors:** Danqing Wang, Fei Ye, Zhou Hao

## Abstract

Antibodies are vital proteins offering robust protection for the human body from pathogens. The development of general protein and antibody-specific pre-trained language models both facilitate antibody prediction tasks. However, few studies comprehensively explore the representation capability of distinct pre-trained language models on different antibody problems. Here, to investigate the problem, we aim to answer the following key questions: (1) How do pre-trained language models perform in antibody tasks with different specificity? (2) How many benefits will the model gain if we introduce the specific biological mechanism to the pre-training process? (3) Do the learned antibody pre-trained representations make sense in real-world antibody problems, like drug discovery and immune process understanding? Previously, no benchmark available largely hindered the study to answer these questions. To facilitate the investigation, we provide an **A**n**T**ibody **U**nderstanding **E**valuation (`ATUE`) benchmark. We comprehensively evaluate the performance of protein pre-trained language models by empirical study along with conclusions and new insights. Our `ATUE` and code is released at https://github.com/dqwang122/EATLM.

## 1 Introduction

Antibodies are special type of proteins extensively used as diagnostic and therapeutic tools against diverse diseases, such as SARS-COV-2 (Zhu et al., 2022). Deciphering the information stored in antibody sequences is highly important with regard to real-world therapeutic antibody development and immune understanding (Greiff et al., 2020; Lu et al., 2018; Yermanos et al., 2018). Fortunately, the recent progress of general **P**re-trained **P**rotein **L**anguage **M**odels (PPLM) and specific **P**re-trained **A**ntibody **L**anguage **M**odels (PALM) provide new prospects for antibody-related tasks. For example, many studies have shown that the representations learned by PPLMs are promising to transfer to antibody tasks (Kim et al., 2021; Zaslavsky et al., 2022). For PALMs, they have been shown to improve model performance in antibody paratope predictions (Leem et al., 2022).

Despite the current success, few studies comprehensively investigate the representation capability of different pre-trained language models (e.g. general PPLM and specific PALM) on distinct antibody tasks, which limits our ability to design better architectures that can help antibody discovery and modification. For example, in Figure 1, we compare the performance of the pre-trained protein language model ESM (Rives et al., 2021), the pre-trained antibody language model AntiBERT (Leem et al., 2021), and the model trained from scratch on three antibody tasks. These three tasks range from low to high in terms of antibody specificity. Here, specificity refers to the antibody’s evolution, for example, the antibody evolves to obtain the ability to bind antigen (The definition is discussed in detail in Section 3.1 and Appendix A.1).

**Figure 1:**
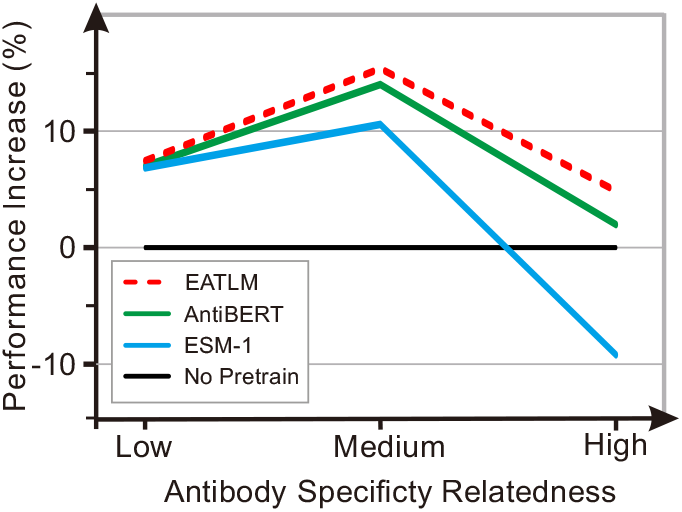
Performance of pre-trained language models on tasks with different specificity. ESM-1 belongs to PPLMs and AntiBERT belongs to PALMs. EATLM is the method with a specific antibody mechanism introduced.

It can be observed that although ESM performs well in task that has low relevance with antibodies, the performance dramatically degrades in tasks of higher relevance. Besides, AntiBERT does not show clear advantages over the non-pretrain model in the high-specificity task. The results point out the weaknesses of current pre-training language models for antibody studies Direct adaptation of general PPLM representations may damage the performance; Pre-training strategies for PALMs cannot fit the specific antibody biological function well. All of these call for a comprehensive guideline of model designing for different antibody tasks. Mainly, we focus on answering the following questions:

### (I) How do pre-trained language models perform in antibody tasks with different specificity?

Addressing of the question is mainly hindered by two challenges: the lack of a reliable antibody-specific benchmark for performance evaluation and a comprehensive study of current PPLMs and PALMs. **(II) How many additional benefits will the model gain if the biological mechanism is introduced to the pre-training process?** Antibody-specific evolution has been employed by many computational biology studies and shows promising results in antibody-related tasks, such as disease diagnosis and therapeutic antibody development (Yermanos et al., 2018; Miho et al., 2019). Then, it is interesting to know whether antibody representation learning can benefit from the incorporation of antibody-specific evolution information. **(III) Does the learned antibody pre-trained representations make sense in real-world antibody problems, like drug discovery and immune process understanding?** Since antibody is so important that is extensively used for drug development in the real world, it is interesting to know whether pre-trained representation can indeed help biologists to understand antibody functions or discover drugs.

To investigate these questions, we first propose antibody study benchmark **A**n**T**ibody **U**nderstanding **E**valuation (**ATUE**). This is the first antibody benchmark with four real-world supervised tasks covering therapeutic antibody engineering, B cell analysis, and antibody discovery. To evaluate models on different aspects of antibody biological functions, these tasks are designed to have specificity ranging from low to high. Based on ATUE, we perform comprehensive empirical studies to investigate the representation ability of distinct pre-trained language models. We also explore the influence of involving specific biological mechanisms in antibody pre-training by introducing two simple objectives to tailor masked language modeling for evolution: (1) *Ancestor germline prediction* guides the model to discriminate the evolutionary relationship between antibody and ancestral sequences. (2) *Mutation position prediction* mimics hypermutation during the evolution. The method is used as an analytic tool to investigate the representation ability of antibody evolution-tailored language model. Finally, we take a close look at the SARS-CoV-2 antibody discovery to investigate the pre-trained representation under a real-world scenario.

Our contributions can be divided three-fold.

- For facilitating antibody application studies, we develop the first comprehensive antibody benchmark to benefit the community. We also introduce two objectives tailored for antibody evolution.
- For providing guidelines for a better antibody representation, we have the following key observations: **(i)** PPLMs perform well on antibody tasks that have a high relationship with structure, such as the binding with antigen. However, they perform worst in tasks with high antibody specificity, indicating representation benefiting protein structure prediction is harmful to antibody-specific tasks. **(ii)** In most cases, PALMs perform as well as or even better than PPLMs with less pretraining data. **(iii)** PALMs can be improved by incorporating the evolution process. However, the evolution information from MSAs does not always benefit the antibody tasks. **(iv)** The introduction of two antibody biological mechanisms facilitates PALMs with more antibody-specific features and improves model performance in the task with high antibody specificity. This is the first attempt showing how antibody specific evolutionary information can be incorporated in pre-training language model.
- For accelerating real-world antibody discovery, we identified 11 potential SARS-CoV-2 binders whose sequences are highly identical to existing therapeutic antibodies binding with the virus.

## 2 Related Work

Our work focuses on researching the effectiveness of protein and antibody pre-trained language models for antibody-specific tasks. Below we review the representative existing methods. We list the details in Table 1.

**Table 1:** Pre-training language models for protein and antibody. Evolution denotes whether evolutionary-related sequences are used during the pretraining. MLM is masked language modeling pretraining objective. HC, antibody heavy chain; LC, antibody light chain.

| Model | Category | Dataset | Evolution | Objective | Antibody Tasks |
| --- | --- | --- | --- | --- | --- |
| ESM-1 (Rives et al., 2021) | PPLM | UniRef50 (27M) | × | MLM | - |
| MSA-1b (Rao et al., 2021) | PPLM | UniRef50 (26M MSAs) | ✓ | MLM | - |
| Ablang-H (Olsen et al., 2022b) | PALM | OAS (14M HC) | × | MLM | Reconstruction |
| Ablang-L (Olsen et al., 2022b) | PALM | OAS (0.19M LC) | × | MLM | Reconstruction |
| AntiBERTa (Leem et al., 2021) | PALM | OAS (72M) | × | MLM | Paratope Prediction |
| EATLM | PALM | OAS (20M) | ✓ | MLM, AGP & MPP | ATUE |

### Pretrained Protein Language Models (PPLMs)

There is an increasing interest in exploring large-scale language models using protein sequences (Rao et al., 2019; Madani et al., 2020; Meier et al., 2021). These models have been shown to achieve state-of-art capacity in predicting protein structure and function. ProtTrans (Elnaggar et al., 2021) and ESM-1b (Rives et al., 2021) take individual protein sequences as input and adopt Transformer language models for pre-training, demonstrating that self-supervision is a promising paradigm for protein secondary structure, contact, homology predictions, and function prediction. To extract evolutionary information from protein sequences, Rao et al. (2021) proposed the MSA-transformer/MSA-1b model utilizing multiple sequence alignment (MSA) instead of a single query sequence as input. This model is superior to ESM-1b for structure prediction, demonstrating evolution information can benefit protein representation learning. Despite the progress in the field, few studies reported protein PLM transfer learning results on antibody tasks.

### Pretrained Antibody Language Models (PALMs)

Encouraged by the success of PLMs in protein representation learning, series work seeks to learn antibody representations based on sequences of antibodies. (Leem et al., 2021; Ruffolo et al., 2021; Olsen et al., 2022b; Prihoda et al., 2022; Li et al., 2022). AntiBERTy (Ruffolo et al., 2021) proposed the first antibody-specific language model, exploring a Transformer trained on 558M natural antibody sequences in the OAS database. The case study in the work showed the representation obtained from the PLM is useful for antibody sequence clustering into trajectories resembling affinity maturation. Olsen et al. (2022b) train two language models for antibodies. A heavy chain version Ablang-H is trained on 14M sequences and a light chain version Ablang-L on 0.19M sequences. The study reported transfer learning results on restoring missing residues of antibody sequences, which is a task similar to pre-training objectives. AntiBERTa (Leem et al., 2021) train the antibody language model on OAS. Although this paper claims finetuning AntiBERTa for paratope position prediction can achieve state-of-the-art performance, the experimental results lack standard deviations, making it unclear how significant the results obtained are. Recently, Li et al. (2022) proposed a antibody-specific language model and explored its performance in SARS-CoV-2 antigen binding, showing context-dependent representations of antibody sequences benefit binding prediction.

## 3 Framework

In this section, we first give a brief introduction to the antibody and its specific evolution. Then we propose the first antibody-specific benchmark (`ATUE`) composed of four tasks with different specificities. Finally, we implement several PPLMs and PALMs baselines and design an evolution-aware PALM to incorporate the biological mechanism into the pre-training process.

### 3.1 Background

#### Antibody

Antibodies are vital proteins generated by the immune system to remove harmful foreign pathogens in the human body. they can specifically bind to antigens on the pathogen and recognize it. Antibodies are composed of two identical heavy chains and two identical light chains and form a large Y-shaped structure. Two tips on it contain highly variable loops, called Complementarity Determining Regions (CDR), which function for antigen binding.

#### Antibody Specific Evolution

Notably, the antibody evolution process is significantly different from that of proteins, providing a good opportunity for us to investigate the impact of general PPLMs on specific subdomains. To perform its protective function, the antibody sequence undergoes evolution selection to search for optimal patterns that can specifically recognize pathogens (Honjo & Habu, 1985). Deciphering the information stored in antibody sequences may benefit our understanding of disease and accelerate therapeutic antibody development (Greiff et al., 2020; Lu et al., 2018; Yermanos et al., 2018). During evolution, the random recombination of V/D/J-gene segments provides the initial diversity for the ancestor sequence (germline). Upon exposure to a pathogen, this sequence undergoes frequent sequence mutations to search for progeny antibody sequences with optimal binding specificity. In other words, gene recombination provides millions of germlines in the human body, and the germlines further mutate into a huge number of progeny antibodies. Thus, the ancestor relationship between an antibody and its corresponding germline as well as the mutation it undergoes together determine the unique biological functions. In brief, the evolutionary relationships between antibodies arise to gain new functions such as antigen binding. It is significantly different from that of proteins, which are to certain functions across different organisms. We further illustrate this process in Figure 7 in Appendix A.1.

#### Unsupervised Antibody Corpus

To obtain the evolutionary information of antibody sequences, we utilize Observed Antibody Space (OAS), a database containing more than 1.5 billion natural antibody sequences (Kovaltsuk et al., 2018; Olsen et al., 2022a) The antibody sequences in the database have been precisely annotated with evolutionary and structural information, including the paired germline and CDR3 for each antibody. To pair the antibody with its germline used in the pretraining task, we used the annotated sequences provided in the OAS database. Further information on data processing can be found in Appendix A.2.

### 3.2 AnTibody Understanding Evaluation (ATUE)

We provide four biologically relevant downstream prediction tasks to serve as antibody benchmarks, covering four major application aspects for antibodies in the real world: therapeutic antibody engineering, disease diagnostics, antibody discovery, and B cell maturation analysis. Notably, the antibody evolution relatedness of these tasks ranges from low to high, offering scaled tasks with subdomain specificity for pre-trained language model evaluation. Detailed information is listed in Figure 2. All data are publicly open and used under the right license. For each task, we focus on the following aspects and leave the details in Appendix:

**[Definition]** The formal definition of the task and the understanding ability required.
**[Impact]** The importance of the task in the biological area. Details in Appendix A.3.
**[Dataset]** The data source and size. Details of data processing in Appendix A.2
**[Specificity]** Antibody’s specific evolution characteristics that is different from general proteins. The definition of antibody specific evolution is shown in Appendix A.1. Quantitative analysis the scale of antibody specificity for every task is shown in Appendix A.4.

**Figure 2:**
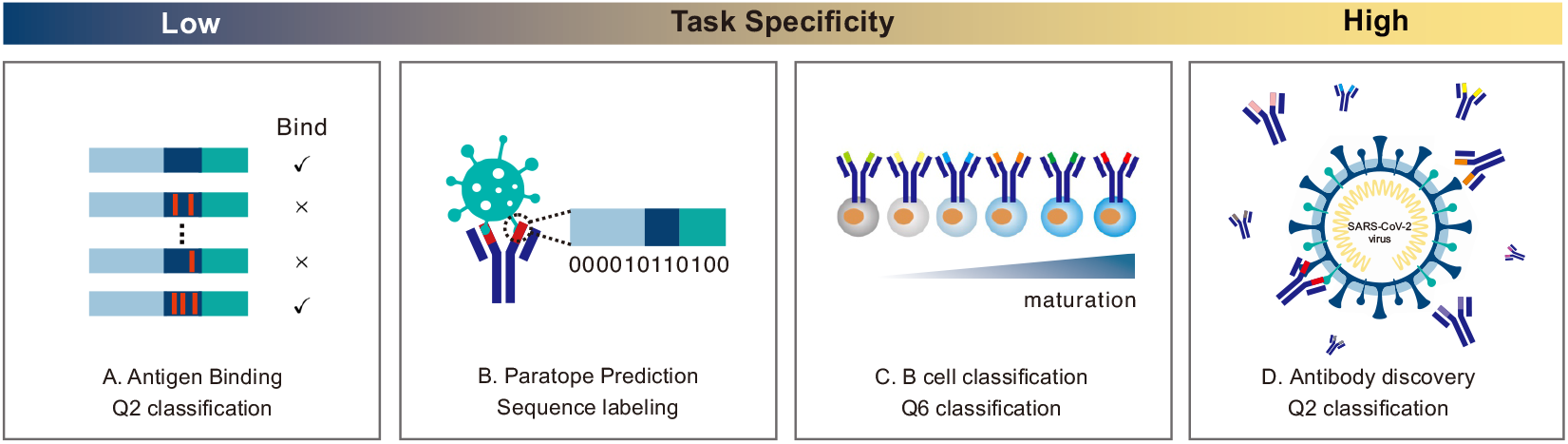
Antibody prediction tasks. The specificity of tasks ranges from low to high.

We use several classification metrics to evaluate the performance. Accuracy (**ACC**) calculates the ratio of correct predictions. Matthews Correlation Coefficient (**MCC**) is the coefficient between true and predicted values. **F1** is the average weighted score of precision and recall. **AUC** is the area under the ROC curve, which shows the performance at all classification thresholds.

**Antigen Binding Prediction** is a binary sequence classification task to determine whether the CDR region of the antibody can bind to the specific antigen.

**[Impact]** A better understanding of the binding affinity between antibody and antigen can accelerate the affinity optimization of therapeutic antibodies.
**[Dataset]** We collect the antigen binding data from (Mason et al., 2021) and follow the training/validation/test split of 15,128/3,242/3,242. The paratope data is collected from (Liberis et al., 2018) with 1,662 CDR segments on 277 antibodies.
**[Specificity]** Low. All the antibodies sequence in the dataset are derived from a single germline sequence indicating the task is not antibody-specific evolution-related.

**Paratope Prediction** is to identify binding positions on the antibody sequence, which is a sequence labeling task to predict a 0/1 label for each residue of CDR fragments.

**[Impact]** The exploration of paratope (binding positions between antibody and antigen) can help to understand the binding mechanisms of therapeutic antibodies.
**[Dataset]** The paratope data is collected from (Liberis et al., 2018) with 1,662 CDR segments on 277 antibodies.
**[Specificity]** This task is medium specificity related because only partial antibodies from the database are derived from evolution.

#### B Cell Maturation Analysis

It is a 6-category classification task to distinguish the maturation stage of B cell antibody sequences. Each sequence belongs to one of {*immature, transitional, mature, plasmacytes, memory IgD*+, *memory IgD*-}. It requires the model to learn a representation sensitive to different maturation states.

**[Impact]** It benefits the understanding of the mechanism during immune evolution, which is a critical biological process in the immune system affecting the function and antigen specificity of antibodies (Ghraichy et al., 2021; Meffre et al., 2000).
**[Dataset]** We collect 88,094 sequences from (Mroczek et al., 2014).
**[Specificity]** High. Antibody evolution is highly coupled with B cell maturation. (Meffre et al., 2000)

#### Antibody Discovery

The task is a binary sequence classification task to distinguish which antibody is directly responsible for SARS-CoV-2 binding. The task is highly challenging from two aspects: (1) Less than 1% of antibodies from SARS-COV-2 patients are directly responsible for virus binding. (2) It is hard to get a reliable sequence-level classifier using unreliable and noisy individual-level labels.

**[Impact]** Antibody discovery from B cell repertoire has been widely recognized as a important approach to accelerate antibody discovery for diverse antigens (Weiner, 2015; Pedrioli & Oxenius, 2021), and achieved great success for SARS-CoV-2 antibody discovery (Kovaltsuk et al., 2018; Cao et al., 2020; Shiakolas et al., 2022).
**[Dataset]** We collected antibody sequences from 133 SARS-CoV-2 patients and 87 health persons from OAS and followed the processing pipeline of (Kim et al., 2021). Inspired Zaslavsky et al. (2022), we match the high-ranked sequences with the sequences in the CoV-AbDab (Raybould et al., 2021) database, which have been proved to bind SARS-CoV-2 using wet-lab experiments.
**[Specificity]** High. It is widely reported antibodies derived from the same disease such as SARS-CoV-2 share strong convergent germline signals (Galson et al., 2020).

### 3.3 Experiment Setup

Based on the antibody benchmark `ATUE`, we evaluate the performance of current pertaining language models in different specificity tasks. Furthermore, to investigate the benefit of introducing the biological mechanism, we incorporate evolution information as the extra pretraining objectives for PALMs and propose `EATLM`. The detailed description can be found in Appendix A.5

#### Current Pre-trained language models

Existing antibody and protein language models are summarized in Table 1. Since the code and pre-training data of AntiBERTa are not released, we train a BERT model named **AntiBERT** on the full OAS database following the same setting as the original study. **MSA-1b** (Rao et al., 2021) takes protein-specific evolutionary sequences (Multiple Sequence Alignment, MSA) as the input. Because it is hard to align sequences between antibodies due to the diversity of CDR3, we take the germline and create pseudo-MSAs with depth 2. We add a linear layer on top of the language models and finetune the whole model on the downstream tasks.

#### Evolution-aware antibody pretraining method

To incorporate the biological mechanism into the pre-training, we propose a model with evolution information: **A**ntibody **E**volu**T**ion-aware pretraining **L**anguage **M**odel. The antibody can be represented as *A* and the germline of the individual antibody can be represented as *G.* Typically, PALMs are trained with basic masked language modeling (MLM). Based on it, we design another two pre-training objectives to simulate the biological mechanism of antibody evolution. The evolutionary relationship between the antibody and its germline includes two folds: (i) *Whether the antibody and the germline have an evolutionary relationship.* (ii) *How to mutate residues from the germline to get the specific antibody.* Two evolution-related objectives are introduced to solve the above questions: **ancestor germline prediction (AGP)** and **mutation position prediction (MPP)**. For ancestor germline prediction, we substitute the paired germline *G* with random germline *G*′ in the batch via a probability *p.* The model is made to distinguish the ancestor germline of the antibody by capturing the shared features. To predict mutation position, for each token in the germline *G*, the objective is to predict a 0/1 label for each token to indicate whether this token has been mutated. For the antibody sequence *S,* we mask the mutation position and predict these tokens. The technical details are described in Appendix A.5.1.

#### Hyper-parameters

We use the base Transformer architecture (Vaswani et al., 2017) with 12 layers, 12 heads, and 768 hidden states. For each task in `ATUE`, we finetune the model with supervised data. We follow the standard split of Antigen Binding Prediction. For other tasks that do not provide a standard split, we use a 10-fold cross-validation. Since our pre-training model learns the representation of the antibody sequence, we expand the CDR fragment to the full antibody by searching the biological database for therapeutic antibody engineering tasks. We also use the same Transformer architecture to train from scratch for each downstream task. This model is indicated as **non-pretrain** since it is not pre-trained on a protein/antibody database.

#### Reproduction

We conduct 10-fold validation on paratope prediction, B cell maturation analysis, and antibody discovery. For antigen binding prediction, we conduct three repetitive experiments with different random seeds. We report the average results and the standard derivation. The benchmark and code will be released.

## 4 Results and Analysis

### 4.1 Supervised antibody prediction

#### Antigen binding

Here, we evaluate the performance PLMs models for antibody binding and paratope prediction, which are less antibody specific. The results are shown in Table 2 It is observed that PPLMs and PALMs achieve comparable performance on this task, indicating PALMs can learn similar general protein representation as PPLMs. Among different PALMs, Ablang-H outperforms Ablang-L and AntiBERT. It indicates that separate training for heavy and light chain sequences is helpful for this task. The introduction of AGP and MPP provides a little improvement over AUC and F1 metrics.

**Table 2:** Performance of PPLMs and PALMs on antibody tasks with increasing specificity. The reported results are the average of repetitive experiments with the standard derivation. `EATLM` w/o AGP indicates that we remove AGP objective from the pre-taining phase.

|  | Antigen Binding (Low) |  |  | Paratope (Medium) |  |  | Cell (High) |
| --- | --- | --- | --- | --- | --- | --- | --- |
|  | AUC | F1 | MCC | AUC | F1 | MCC | ACC |
| non-pretrain | 0.858±0.014 | 0.584±0.330 | 0.432±0.183 | 0.845±0.014 | 0.605±0.033 | 0.463±0.037 | 0.554 ± 0.042 |
| ESM-1 | 0.917±0.001 | 0.854±0.002 | 0.689±0.002 | 0.886±0.009 | 0.669±0.024 | 0.547±0.026 | 0.503 ± 0.031 |
| MSA-1b | 0.921±0.001 | 0.857±0.004 | 0.689±0.014 | 0.887±0.009 | 0.679±0.019 | 0.557±0.025 | 0.416 ± 0.050 |
| Ablang-H | 0.918±0.001 | 0.861±0.003 | **0.704±0.010** | 0.878±0.009 | 0.674±0.018 | 0.546±0.023 | 0.570 ± 0.010 |
| Ablang-L | 0.917±0.002 | 0.856±0.001 | 0.682±0.001 | 0.882±0.010 | 0.680±0.018 | 0.553±0.023 | 0.546 ± 0.008 |
| AntiBERT | 0.918±0.003 | 0.843±0.009 | 0.678±0.008 | 0.879±0.011 | 0.690±0.020 | 0.559±0.026 | 0.565 ± 0.028 |
| EATLM | 0.922±0.004 | **0.862±0.004** | 0.699±0.010 | **0.887±0.008** | **0.698±0.017** | **0.575±0.024** | **0.581 ± 0.005** |
| EATLM w/o AGP | 0.920±0.003 | 0.855±0.000 | 0.697±0.008 | 0.883±0.011 | 0.676±0.021 | 0.552±0.027 | 0.559 ± 0.012 |
| EATLM w/o MPP | **0.923±0.001** | 0.855±0.002 | 0.687±0.005 | 0.883±0.011 | 0.691±0.022 | 0.566±0.030 | 0.563 ± 0.010 |
| EATLM w/o AGP & MPP | 0.918±0.001 | 0.845±0.005 | 0.681±0.005 | 0.880±0.009 | 0.674±0.018 | 0.552±0.023 | 0.559 ± 0.009 |

#### Paratope prediction

As the results shown in Table 2, for paratope prediction, both PPLMs and PALMs can significantly boost the prediction accuracy over the model with pre-training. However, PALM doesn’t show much significant advantage over PPLMs. `EATLM` achieves the best performance, especially on F1 and MCC, while other pertaining models have a high recall and a low precision. It indicates that those models tend to predict more residues as the binding site. However, with the introduction of mutation residue prediction, `EATLM` can focus on the mutated positions that are adapted to bind with antigen. Among the two PPLMs, MSA-1b outperforms ESM-1 on F1 and MCC, which benefits from the structure information learning from MSAs.

#### B Cell Analysis

In this task, we examine the representation capacity of different pre-trained language models for distinguishing different B cell mature states during evolution. The results are shown in Table 2. In general, we find PPLMs can hardly distinguish the minor differences between B cell sequences by showing compromising results. Both the performance of ESM-1 and MSA-1b is significantly worse than randomly initialized models. MSA-1b performs the worst in all pre-trained language models, indicating representation that can perform well in protein structure prediction will be harmful to antibody-specific tasks. Conversely, all PALMs show promising results for the task. The reason may lie in that the general protein has little relationship with the special antibody mature process and can hardly capture this feature during the protein pretraining process. `EATLM` significantly outperforms the other PALMs. This is because by explicitly modeling the biological mechanism, our model can effectively capture the evolution feature and better distinguish between B cells at different stages of maturation.

We conduct further analysis to figure out whether our `EATLM` successfully captures sequence characteristics during the evolutionary process. We explore the probabilities of predicting antibodies in class *i* to class *j*. The results shown in Figure 3 reveal `EATLM` can easily classify the immature B cell with an accuracy of 0.9. It is consistent with the biological study that CDR3 sequence length in immature B cells is significantly shorter than that of the other mature B cells (Ghraichy et al., 2021). From the diagonal, we can figure out that our model tends to mistake the B cell sequences with their previous or post-evolutionary stage, consistent with the biological process.

**Figure 3:**
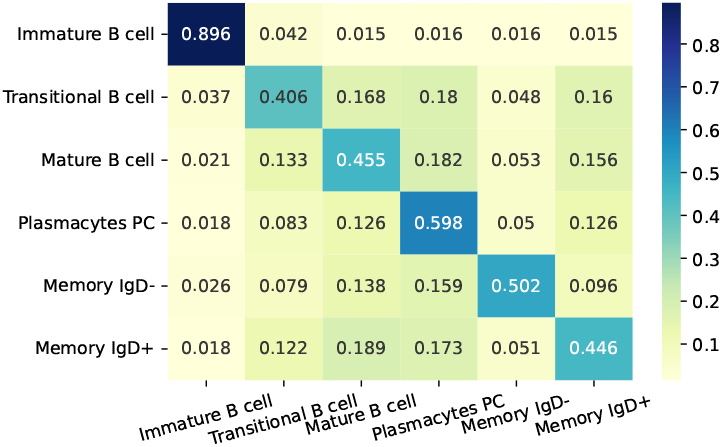
B cell evolution category prediction. For each *p_ij_* in *i*–th row and *j*-th column, it means the frequency for the model to predict the antibody in *i* category to *j* category. The number is normalized by row.

#### Antibody Discovery

Here, we examine PPLMs and PALMS for their capacity in benefiting real-world problems and enabling the discovery of antigen-specific antibodies. Following the methods in this study Zaslavsky et al. (2022), we considered two steps for the SARS-CoV-2 specific antibody discovery. First, we generate a sequence classifier to distinguish SARS-CoV-2 antibodies using noisy individual-level labels. Then, the high-ranked sequences are matched with the sequences with true binding sequences in the CoV-AbDab (Raybould et al., 2021) database. We use the 90% sequence identity as the threshold to determine whether the antibody is similar to existing SARS binders in the CoV-AbDab database. A high similarity indicates a high probability of having the same biological functionality as the existing binders. The details can be found in Appendix A.7.

In Figure 4, we plot the cumulative sum of matched sequences number in the order of the predicted probability of the classifiers. We can find that the sequences predicted with high probability by PALMs match with the existing binders better than PPLMs. It means that PALMs can figure out the potential binders more efficiently. Besides, `EATLM` significantly outperforms other PALMs as the red line shows. In the early stage, this method is the quickest way to find potential binders. Then it loses to Ablang-H but finally overtakes again and converges. It means that `EATLM` is the first one to figure out all potential binders in this dataset.

**Figure 4:**
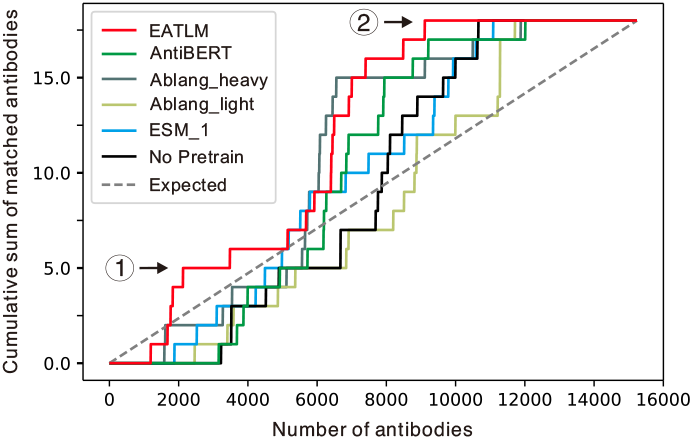
The cumulative sum of matched sequences number in the order of the predicted probability. EATLM outperforms other PALMs and PPLMs for finding SARS-CoV-2 binder faster highlighted in ① and finding all antibodies faster highlighted in ②.

Furthermore, we list several potential binders found by `EATLM` in Table 3. Without supervised labels, `EATLM` gives a high probability of 2 SARS-CoV-2 existing binding antibodies. Besides, `EATLM` proposes 9 potential sequences with high CDR-H3 sequence identity. Further analysis of the results reveals that `EATLM` enables diverse-epitope antibody discovery and priority selection, demonstrating that `EATLM` benefits therapeutic antibody discovery. The detailed explanations can be found in Appendix A.7.

**Table 3:**
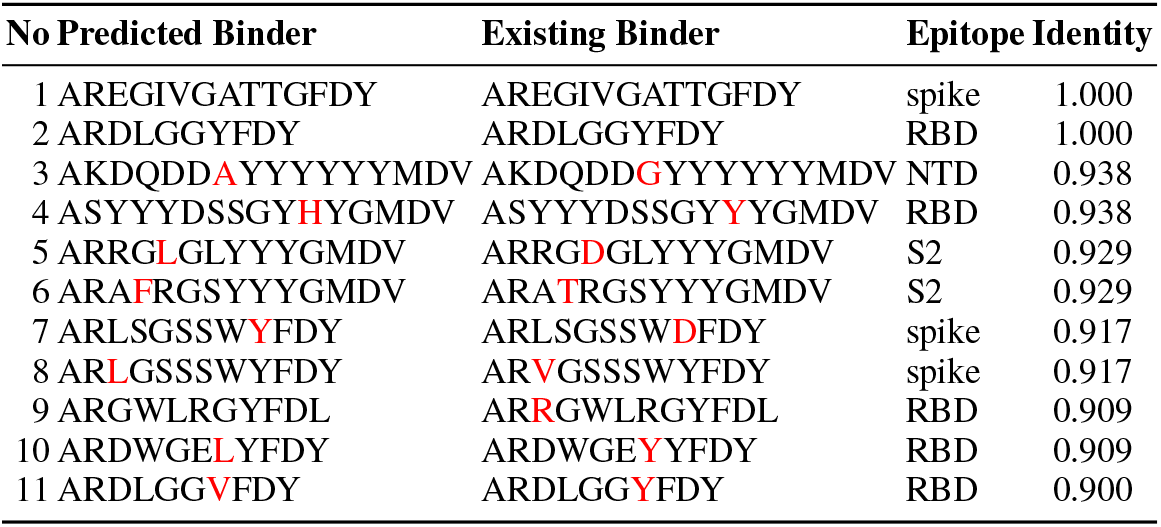
The CDR3-H3 region of high-ranked sequences to bind to SARS. We show the CDR3 fragment of the heavy chain in the antibody sequences. ‘Identity’ is the similarity between the predicted binder and the true binder. The epitope of the true binders is shown. The origin of the majority of the true binder sequences is B cells from patients. The different amino acids between the predicted binder and the existing binder are highlighted in red

To examine whether the antibody sequences with 90% sequence identity can indeed bind the same target, we investigate the 3D structure of the true binding antibody. Table 4 shows only one signle atom difference between the predicted binder and the existing binder, suggesting the predicted binders are highly possible to interact with SARS-CoV-2.

**Table 4:**
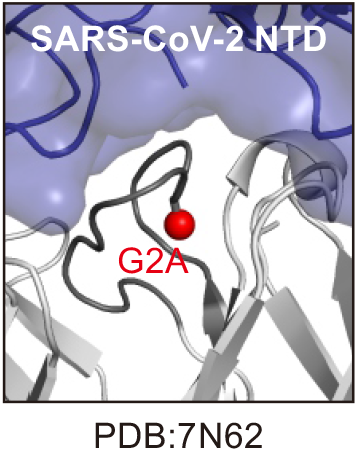
3D structure of the true SARS-CoV-2 binding antibody No.3. G2A highlights the single atom difference in No.3, indicating the predicted binder is highly likely to bind the virus.

### 4.2 How does evolution pretraining task influence the representation?

To understand why `EATLM` shows better performance on antibody tasks, we analyze the pre-trained representations to evaluate the effectiveness of the evolution-aware pretraining strategies from two aspects: (1) Does the pre-trained antibody representation reflect its ancestor relationship? (2) Is the specificity of antibodies captured by the evolution objective?

#### Ancestor Gerlime Visualization

We perform UMAP visualization analyses in Figure 5. First, we observe that antibodies evolved from the same germline are nicely clustered together (Figure 5a and 5b), indicating the learned embedding is encoded with germline information. Besides, sequences with similar scales of evolutionary distance tend to cluster together, and a clear gradation of evolutionary distance can be observed in Figure 5c and 5d. The visualization provides a sanity check for the ability of EATLM to extract the sequence information of antibody.

**Figure 5:**
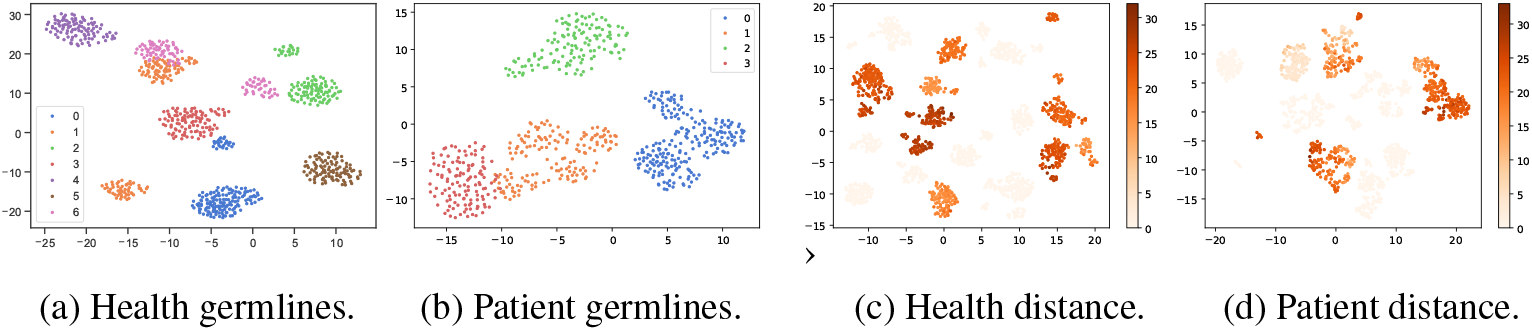
UMAP Visualization. (a-b) for sequences with different germlines. One color indicates one germline with its descendant. (c-d) for evolutionary phylogeny distance. The shade of color indicates its distance from the ancestor germline.

**Figure 6:**
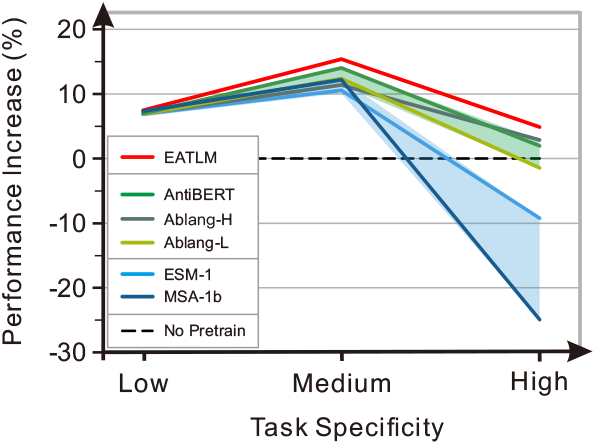
Performance summary of various pre-trained language models.

#### Accuracy of Mutation Position

Based on the specific evolution process described in Section 3.1, we can find the mutation during the evolution process bring specificity to the antibody. Thus, we explore the model’s ability to predict mutated residue from the masked token, which can reflect the specificity feature the model captures. We find that although AntiBERT can predict with an accuracy of 0.8889 on all positions, it fails on mutation positions with a 0.0311 accuracy. In contrast, `EATLM` achieves an accuracy of 0.443 on mutation position, which indicates that the model captures the specificity information. Note that during the MPP training, we mask the mutation position on antibody sequences, which are different from its germline. Thus, the model cannot get the mutated residue from the germline directly. The only way is to learn the underlying mutation rules. The full results are shown in Table 12a in Appendix.

### 4.3 Key Observations

#### The performance of pre-trained language models highly depends on the task specificity

For tasks with low antibody-specificity relatedness, PPLMs show comparable performance with PALMs, indicating that transferring the general protein representations from PPLMs is an effective and time-saving way in these tasks. PALMs show their advantage and outperform PPLMs on medium specificity relatedness tasks, like paratope prediction. For tasks with high specificity, it is observed the performance of PPLMs dramatically decreased, suggesting general pre-trained protein models are not sufficient in antibody-specific representation learning. Interestingly, the performance of MSA-1b is 20% less than the model without pre-training. This observation is consistent with the biological study that the mechanism of antibody evolution is significantly different from that of proteins. Therefore, the incorporation of protein evolution information is not always good for antibody tasks, especially the tasks that require antibody evolution information.

#### Incorporation of biological evolution mechanism into PALM generally benefits antibody prediction tasks

Evolution-related training objectives help detect mutation positions on antibodies, which is a distinguishing feature from germline. Interestingly, the performance increase of `EATLM` compared with other PALMs is correlated with task specificity relatedness. The ablation study showed the removal of the evolution-related pretraining objectives results in performance decreasing, confirming that evolution-related objectives benefit the prediction. Future studies to have more in-depth research in this direction could be promising.

#### Antibody pre-trained representations are helpful for real-world drug discovery

Using the language model, we predict the probability for each antibody of being a binder with SARS-CoV-2. Without accurate sequence-level labels, we successfully identify 11 potential antibody binders.

## 5 Conclusions and Limitations

In this paper, we have presented comprehensive empirical studies to characterize the impact of pre-trained protein and antibody language models for antibody prediction tasks. Four important antibody tasks from four biological categories with evolution relatedness ranging from high to low are provided `ATUE` to facilitate the antibody and ML field.

However, there are some limitations to this work. First, for the `ATUE`, the task diversity is highly limited because of data scarcity. With the data increase in this field, we expect we can update our benchmark with more disease diversity and data size in the future. Also, we do not test any 3D structure information during antibody pre-training. As a special subgroup of proteins, antibody structures provide much more information such as geometry than sequences. In the future, recruiting structure information for antibody pre-training may be able to improve the results.

## A Appendix

### A.1 Antibody Specific Evolution

Antibodies, composed of two identical heavy chains and two identical light chains, form a large Y-shaped structure, where the two tips are responsible for pathogens binding. Antibody evolution, described by sequence-sequence relationships between ancestor and progeny antibodies, reflects antibodies’ key antigen-binding function (Honjo & Habu, 1985). During antibody evolution (Figure 7), the initial diversity is encoded into the ancestor sequence through randomly recombination of V-, D- and J-gene segments. Upon exposure to a pathogen, the sequence undergoes frequent sequence mutations to search for progeny sequences with optimal binding specificity. Sequence evolution analysis has been employed by many computational biology studies and shows promising results in antibody-related tasks, such as disease diagnosis and therapeutic antibody development (Yermanos et al., 2018; Miho et al., 2019).

Importantly, antibody evolution is significantly different from that of proteins. Antibodies only contain hundreds of thousands ancestor sequences so-called germline. To bind dozens of millions of diverse antigens, antibodies need to mutate from the ancestor sequences to gain new functions (Figure 7). Therefore, the **non-conserved amino acids** (mutated ones) plays important roles for structure and function. On the contrary, the **conserved amino acids** (not mutated) in proteins determine structure and function. During protein evolution, evolutionary pressure to maintain protein structure and functions leads to the conservation or co-evolution of residues located in structural folding core for binding interface. Diverse methods have been developed to extract the co-evolution information from conserved amino acids sequences for structure and function prediction, such as AlphaFold (Jumper et al., 2021).

In brief (Figure 7), antibody evolution specificity distinct from that of proteins can be defined with two main features: (i) ancestor germlines; (ii) the mutated amino acids of germlines.

**Figure 7:**
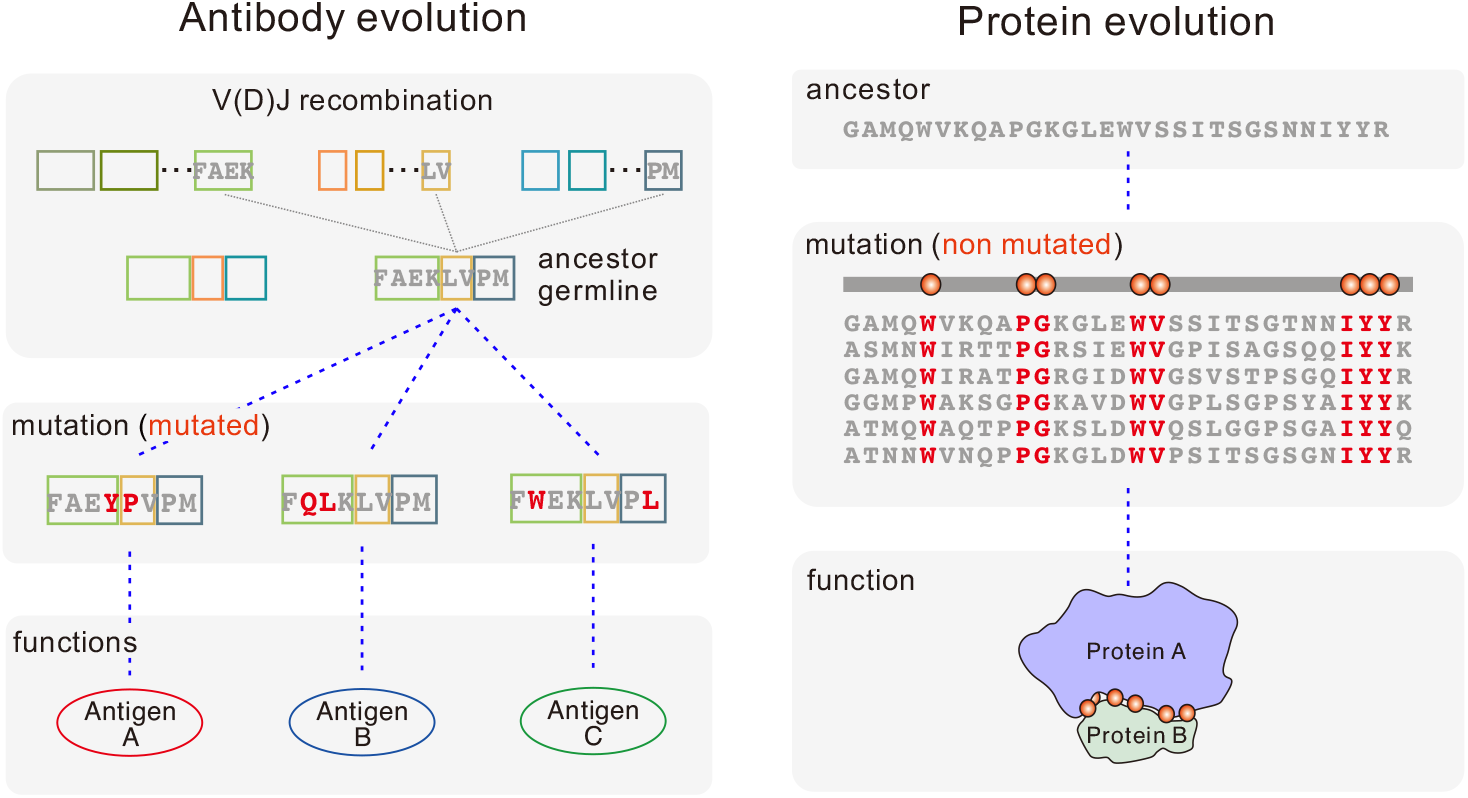
Evolution specificity of B cell antibodies comparing with general proteins. The evolutionary sequence relationships are highlighted using blue dash lines.

### A.2 Data Processing Details

#### Pairing Antibody with Germline

For germline annotation in the pre-training task, we used the annotated germline sequences provided in the OAS database (Kovaltsuk et al., 2018). For downstream benchmarks tasks like B-cell classification, therapeutic antibody engineering, and disease diagnosis, we completely followed the methods shown in the OAS database paper. IgBLAST, an immunoinformatic benchmarking tool for the analysis of B-cell antibody repertoires was used for germline annotation (Ye et al., 2013). The antibody nucleotide-containing FASTA file was aligned to germline and translated to amino acids using IgBLASTn. The antibody amino-acid sequence was aligned using IgBLASTp. The germline databases for human patients used ImMunoGeneTics (IMGT) germline sequences derived from IMGT (Lefranc et al., 1999). For each antibody, usually, multiple germline sequences can be obtained and only the single sequence showing the highest confidence score for the alignment was chosen.

#### Pre-training Data Processing

We downloaded OAS Oct 2021 version from its website and removed duplicate sequences. To avoid data leakage, we cluster sequences based on the CDR3 sequence and filter each cluster by 70% identity over the whole sequence using Linclust (Steinegger & Söding, 2018). Then, we shuffle the dataset and split it into 100k-size chunks. The last chunk is used as the validation set. The dataset size is 20,245,249 and 45,249 are used for validation.

### A.3 ATUE Details

We summarize the tasks used in `ATUE` in Table 5 and discuss each task in detail in this section.

**Table 5:** `ATUE` benchmark. The five downstream tasks are divided into three biological categories and each category focuses on different input levels. ‘Q6’ indicates the classification task have 6 classes. For disease diagnosis, the size indicate the number of profiles, where each profile contains thousands to millions of sequences.

| Specificity | Task Name | Input | Formalization | Size |
| --- | --- | --- | --- | --- |
| Low | Antigen Binding Prediction | CDR fragment | Q2 Cls. | 21,612 seqs |
| Medium | Paratope Prediction | CDR residue | Q2 Labeling | 1,662 seqs, 21,342 positions |
| High | SARS Antibody Classification | Sequence | Q2 Cls. | 22,000 seqs |
| High | B Cell Classification | Sequence | Q6 Cls. | 88,094 seqs |

#### Antigen Binding

Accurate antigen-binding prediction approaches could allow significantly more efficient antibody discovery with higher affinity. Machine learning methods have already achieved some success in antibody binding capacity optimization. We collect the antigen-binding data from Mason et al. (2021) and follow the training/validation/test split of 15,128/3,242/3,242. The original dataset only has CDR3 fragments, and we extend them to the full antibody sequences. For cross-validation, we split the dataset by antibody sequences to ensure that no antibody sequences overlap between 90% training and 10% validation.

#### Paratope Prediction

Paratope is the antibody residues involved in antigen binding. The ability to accurately map the paratope can provide detailed knowledge about the binding mechanism and accelerate antibody discovery. 1D sequence-based deep learning methods have been employed for paratope prediction. The paratope data is collected from Liberis et al. (2018) with 1,662 CDR segments on 277 antibodies. Each antibody contains three CDR fragments (CDR1, CDR2 and CDR3) in the heavy chain and three CDR fragments in the light chain. We also search the full sequence for each antibody and use the whole sequence as input. For cross-validation, we split the dataset by antibody sequences to ensure that no antibody sequences overlap between 90% training and 10% validation.

#### B Cell Analysis

We formulate a 6-category classification task for B cell maturation analysis, which includes *{immature, transitional, mature, memory IgD+, memory IgD-, plasmacytes,*}. The analysis of B cell maturation plays an important role in understanding the mechanisms underlying B cell responses in immune system Ghraichy et al. (2021); Meffre et al. (2000).

The order of B cell type follows the evolutionary process in the immune system, from immature state to transitional state and finally becomes a memory B cell. Both memory IgD- and IgD+ belong to memory B cells with different isotypes, and they have a high affinity to foreign antigens. Among the other categories, the Plasmacytes PC sequences also have some affinity ability. It is widely reported that changes in antibody sequence patterns correlate with B-cell maturation. Therefore, we use this task to evaluate the representation learning capacity of the language model.

We collect 88,094 sequences from Mroczek et al. (2014). They extracted from the peripheral blood of healthy adults and got six types of B cells with different maturity and antibody sequences. The distribution of various types of B cells in the dataset is shown in Table 6

**Table 6:** The statistics of B cell classification.

| Type | Size |
| --- | --- |
| Immature b cell | 14,145 |
| Transitional b cell | 13,197 |
| Mature b cell | 16,139 |
| Plasmacytes PC | 22,236 |
| Memory IgD- | 8,437 |
| Memory IgD+ | 13,940 |

#### Antibody Discovery

Antibody discovery from B cell repertoire has been widely recognized as a novel trend to improve the efficiency of antibody discovery for diverse antigens (Weiner, 2015; Pedrioli & Oxenius, 2021). However, previous studies highly rely on expensive wet-lab experiments (Cao et al., 2020; Shiakolas et al., 2022). Deep learning based method have shown potential capacity in help antibody discovery by reduce cost and increase efficiency (Widrich et al., 2020; Wang et al., 2022). Here, we ask whether pretrained models can benefit real-world problems and enable fast-track neutralization SARS-CoV-2 antibody discovery.

In the first step, we develop a sequence classifier to distinguish which antibody sequence from the numerous sequences is responsible for the recognition of the SARS-CoV-2. This task is highly challenging since we can hardly get the sequence-level disease label that indicates whether the antibody sequence is related to the disease. Thus, we follow the practice of Roskin et al. (2020); Zaslavsky et al. (2022) to use the individual label as the rough sequence label and train a sequence-level predictor. Then, with the help of a sequence-level predictor, we can give each sequence a most likely label to help antibody discover, whose accuracy has been verified by the excellent results on individual prediction, which may accelerate the discovery of new antibody sequences.

We follow the condition of Kim et al. (2021) to filter SARS-CoV-2 antibody data from the OAS database. The basic condition is *‘Chain = heavy; Isotype = IGHG; BSource = PBMC; Species = human; Vaccine = None’*. We further add the condition of *‘Unique Sequences >= 10000’*. For health/SARS we set the *‘Disease’* field to *‘None’*, *‘ SARS-CoV-2’.* Then we obtain 87/133 patient profiles for each type. To make a balanced dataset, we limit the size of the health profile and mix up the healthy ones and the ones with the SARS-CoV-2. For cross-validation, we randomly split the dataset by profiles 10 times: 90% for training and 10% for validation. We further select sequences with top100 redundancy to make the positive labels more accurate.

### A.4 Quantitative Analysis of ATUE Task Specificity

It is important to include statistical significance tests relative to the antibody-specific features in antibody functional tasks we proposed in the ATUE benchmark. According to the evolution process shown in Figure 7, antibody evolution specificity distinct from that of proteins can be defined with two main features: (i) ancestor germlines; (ii) the mutated amino acids of germlines. We implemented statistical significance tests of (i) ancestor germlines subtype usage; (ii) the number of mutated amino acids in antibodies against different labels of downstream tasks in ATUE to quantitatively assess the “Task specificity”. The analysis is now summarized in Table 7 In Appendix A.1. Generally, it is clearly shown that ATUE benchmark comprises antibody tasks showing different scales of antibody specificity for later on modeling analysis. Moreover, the which are used for statistical analysis of task specificity and pre-training model objectives in our study.

**Table 7:** Task specificity. Summary of the statistical significance test of two antibody-specific features for different tasks in the `ATUE` benchmark.

| Task | Statistical significance (p-value) |  | General Specificity |
| --- | --- | --- | --- |
|  | Germline Usage | Mutations Numbers |  |
| Antigen Binding | Nan | Nan | Low |
| Paratope Prediction | 0.296 | 0 | Medium |
| B cell classification | 0 | 0 | High |
| SARS antibody discovery | 0 | 0 | High |

#### Antigen Binding

In the Antigen binding dataset, both antibody binding and none antigen binding sequences share the same germline subtype sequence (IGHV3.1) (Figure 8A) as well as the same number of germline mutations 8B). Therefore, None of two antibody specific features show significant distribution differences between data with different labels, demonstrating antigen binding is a task with low antibody specificity.

**Figure 8:**
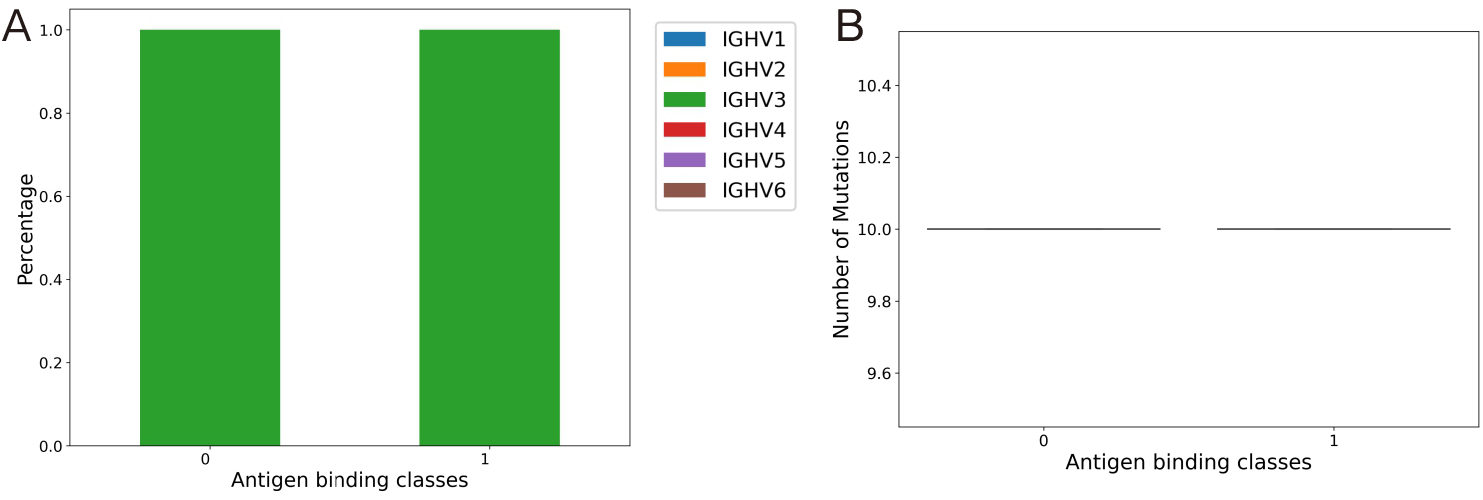
(A)IGHV gene segment usage distribution between binding and non-binding antibody. (B) Germline mutation number distribution between binding antibody and non-binding antibodies.

#### Paratope Prediction

For the paratope prediction task, we first evaluate the germline subtype distribution difference between sequences with different numbers of binding sites (Figure 9A). Kruskal Wallis test showed a p-value of 0.296 suggesting germline subtype usage is not statistically significant. Also, we find the binding sites can be significantly mapped with more germline mutations than the non-binding sites, which is consistent with the knowledge of antibody specificity definition (Figure 9B). One out of two antibody specific features show significant distribution differences between data with different labels. Therefore, we define this task as a medium specificity task.

**Figure 9:**
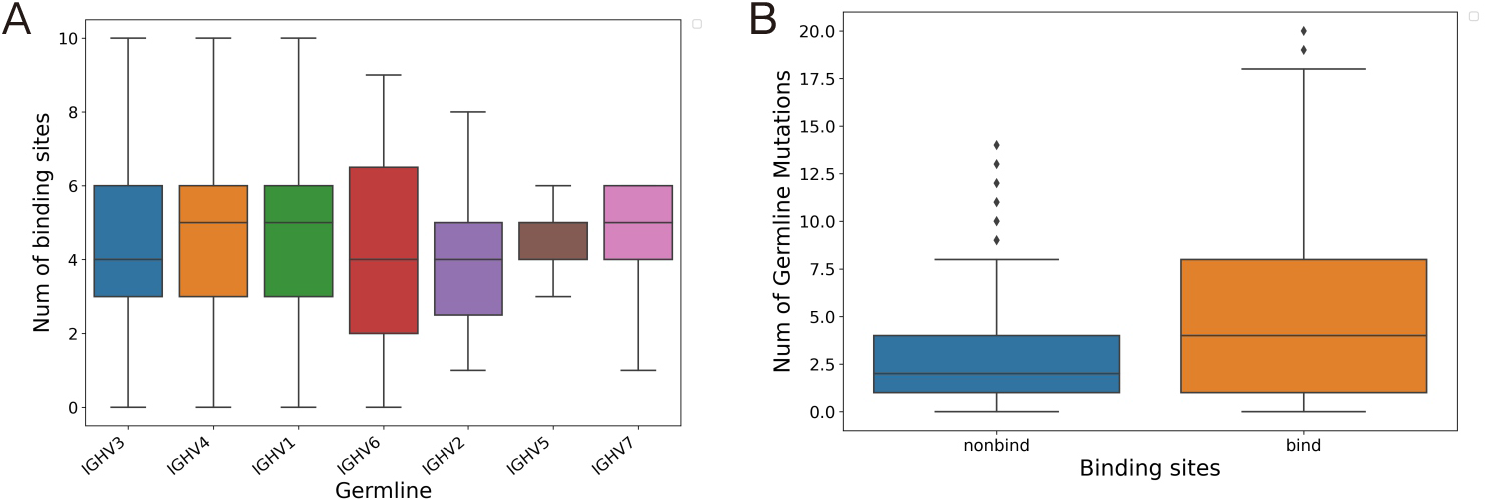
(A) Number of binding sites distribution between different IGHV gene segments. Comparison is performend using kruskal wallis test with p value 0.296. (B) Germline mutation number distribution between binding and non-binding positions. Comparisons performed using t-tests showing p value equals to 0.

#### B Cell Analysis

As shown in Figure 10, the distribution of the germline usage as well as the number of germline mutations are significantly different between antibodies in B cells with different developmental stages. This observation is highly consistent with previous studies Mroczek et al. (2014); Ghraichy et al. (2021). Since both of the antibody specific feature show significant distribution differences, this task is defined as a high-specificity task.

**Figure 10:**
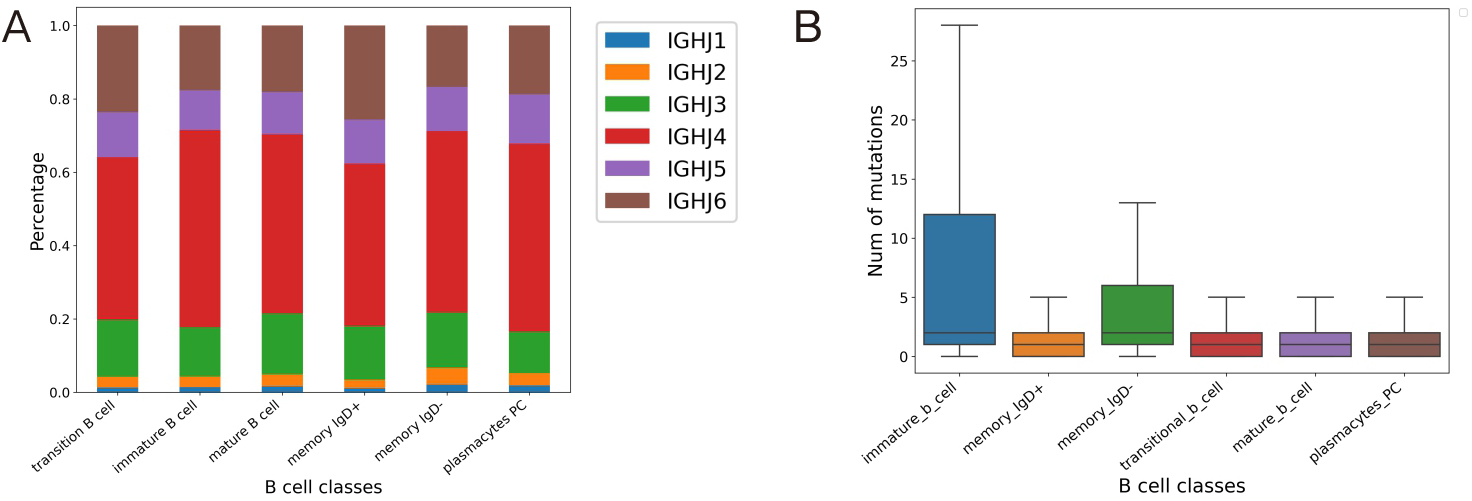
(A)IGHV gene segment usage distribution between different B cells. Comparison is performend using chisquare test with p value 0. (B) Germline mutation number distribution between different types of B cells. Comparisons performed using kruskal wallis test showing p value equals to 0.

#### SARS Antibody Discovery

Antibodies in SARS patients and healthy ones show a significant difference in their germline subtype usage and the number of germline mutations (Figure 11). This observation is highly consistent with previous studies showing SARS antibody convergent among patients Galson et al. (2020). Since both of the antibody specific feature are highly significant, this task is defined as a high-specificity task.

**Figure 11:**
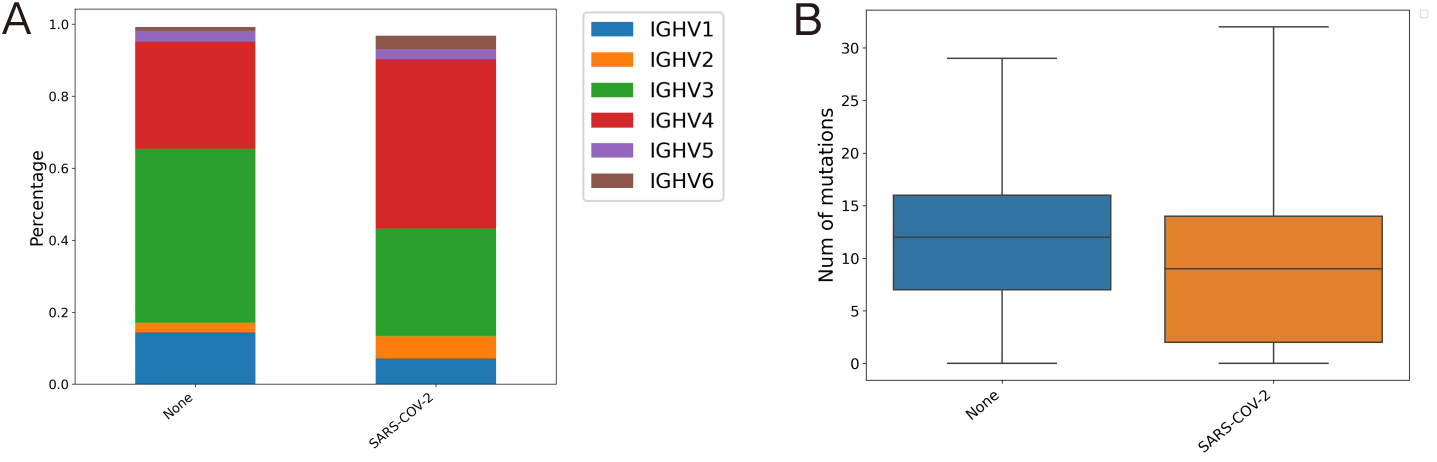
(A)IGHV gene segment usage distribution between antibody in SARS patients and health ones. Comparison is performed using chisquare test with p value 0. (B) Germline mutation number distribution between antibody in SARS patients and health ones. Comparisons performed using kruskal wallis test showing p value equals to 0.

### A.5 Training Details

Antibody can be represented as *A* = {*a*_1_, *a*_2_, ⋯, *a_m_*} and the germline of individual antibody can be represented as *G* = {*g*_1_, *g*_2_, ⋯, *g_n_*}, where *m* and *n* are the lengths. Each token *a_i_* or *g_j_* in the sequence is called a residue that belongs to the amino acid set 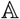. 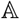 includes 20 common amino acids with a residue ‘X’ that indicates the residue is unknown (mostly in the germline). Typically, antibody PLMs are trained with basic mask language modeling objective *l*_MLM_ on the antibody sequences *S* = *A* { *a*_1_, ⋯, *a_m_*}.

#### A.5.1 Evolution-aware Pretraining

In order to incorporate the evolutionary information into the pre-training, we pair the antibody sequence *A* with its germline *G* and concatenate them into a long sequence with a special token ‘[SEP]’ as the delimiter: *S* = {*s*_1_,⋯, *s*_*m*+*n*+1_} = {*a*_1_,⋯, *a_m_*, [SEP], *g*_1_,⋯, *g_n_*}. Thus, we optimize MLM objective on the long sequence *S*:

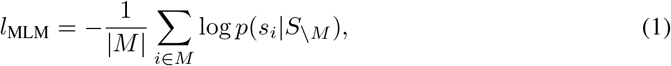

where *M* is the index set of masked tokens. It helps the model to learn the basic residue distribution for antibody sequences. Besides, it can also capture the interaction between residues of the antibody and its germline.

**Figure 12:**
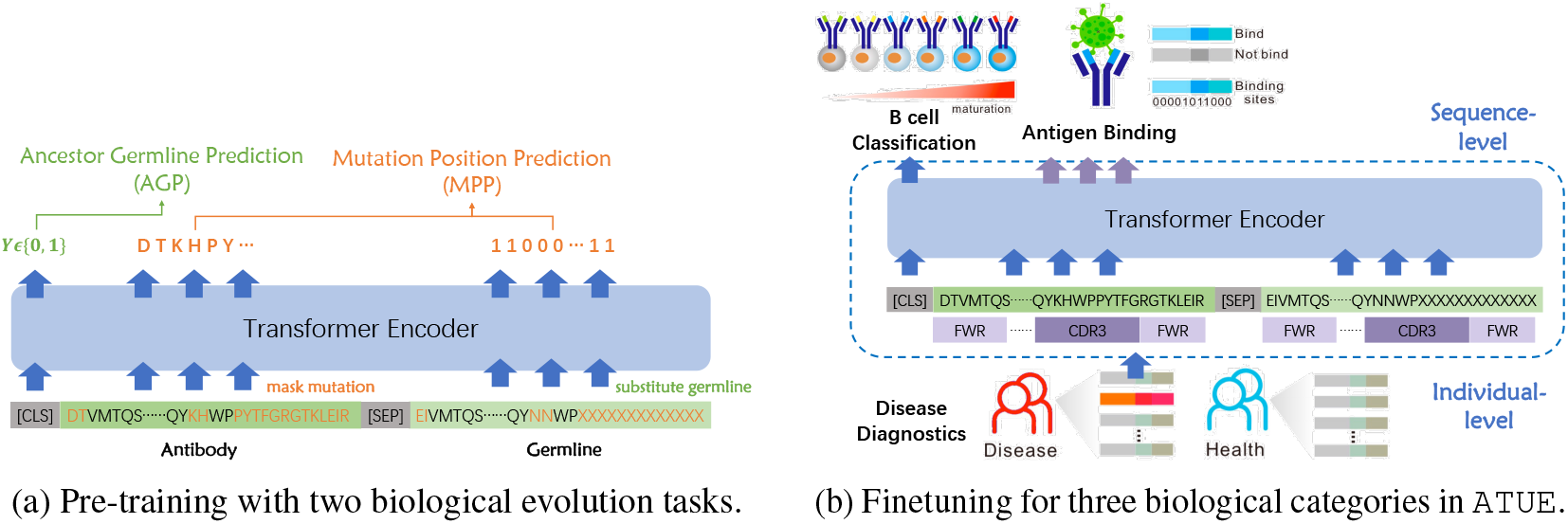
EATLM. In Figure 12a, AGP randomly unpairs the germline sentence and predict the ancestor relationship. MPP predicts the mutation position on the germline and the masked mutation residue on the antibody. Based on the input, the three categories in `ATUE` can be divided into sequence-level and individual-level (Figure 12b). For individual-level disease diagnostics, we score each sequence in the individual profile and calculate the trimmed mean over all sequences to get the individual score.

##### Ancestor Germline Prediction

The ancestor relationship between the antibody and its germline determines the shared biological functions obtained in the evolution. Antibody sequences with similar residues evolved from different germline sequences may have different biological functions. When stimulated by a foreign antigen, the common ancestor germline evolve to various antibody sequences. Similar antibody sequences may have different germline sequences, which will affect their biological functions. Thus, the aim of this task is to determine whether the antibody have evolutionary relationship with the given germline. During training, we substitute the paired germline *G* with random germline *G*′ = {*g*_1_,⋯, *g_n_*} in the batch via a probability *p* = 0.3. The new sequence is denoted as 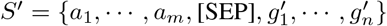 and the training loss can be described as:

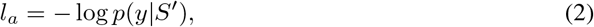

where *y* ∈ {0,1} indicate whether the noisy germline *G*′ is the ancestor of the antibody *S*. It can help the model to distinguish the ancestor germline of the antibody by capturing the shared features.

##### Mutation Position Prediction

The somatic hypermutations on the germline further give progeny antibodies the specificity of binding with the specific antigen. In order to model this specificity, this task focus on predicting the mutation positions and the residues mutated to. Specifically, for each token *g_j_* in the germline *G*, the target is to predict a label *y_j_* ∈ {0,1} to indicate whether this token has been mutated. For the antibody sequence *S*, we mask the mutation position and predict these tokens. The objective can be formalized as:

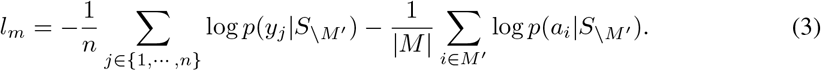

Here, *M*′ is the ground-truth mutation position and we mask these tokens on the antibody sequence. This task is more difficult than MLM which equally masks tokens in the *L*, because the tokens on the mutation position of *A* get less information from the germline, compared with other shared residues between the antibody and the germline. By optimize this objective, the model learn to capture the specificity obtained from the somatic hypermutation in the evolutionary process.

#### A.5.2 Implementation Details

We use the base Transformer architecture (Vaswani et al., 2017) with 12 layers, 12 heads, and 768 hidden states. The total parameters are 86M. We use Adam optimizer (Kingma & Ba, 2015) with the maximum learning rate of 2e-4 and the warm-up step of 24,000. The maximum length is set to 400 since most antibody sequences are shorter than 180. We first pre-train our model with the MLM objective. During the pre-training, 15% tokens are randomly selected with 80% masked, 10% replaced, and 10% kept. Then we conduct further pre-training on two antibody-related tasks with a smaller learning rate of 1e-5.

For each task in `ATUE`, we finetune the model with supervised data. We follow the standard split of Antigen Binding Prediction. For other tasks that do not provide a standard split, we conduct 10-cross validation and report the average results. Since our pre-training model learns the representation of the antibody sequence, we expand the CDR fragment to the full antibody via searching the biological database for therapeutic antibody engineering tasks. For finetuning, we limit the max epochs to 30 and use the Adam optimizer with a max learning rate of 3e-5. We use the mean representation of 12 layers as the sequence representation.

The model is trained with 108,000 steps and gets a 0.9606 token accuracy on the MLM task. It takes further steps for AGP and MPP. The model quickly converges for AGP and gets a 0.99 accuracy on the ancestor germline prediction because more than 80% residues are shared between the antibody and its germline. For MPP, it can predict the mutation position with the accuracy of 1.00 and obtains a 0.4430 accuracy in the mutation position. It means that the model can easily find the mutation positions by the self-attention between the antibody and germline, but it is still difficult to predict which residues this position will mutate to. We assume it is because the ancestor germline can undergo different somatic hypermutations and get various progeny antibodies, resulting in different valid mutations at the same position. We also compare this mutation accuracy with the model without MPP, which is only trained with MLM on the concatenation of the antibody and its germline. With a high prediction accuracy of 0.8889 on all positions, it achieves only a 0.0311 accuracy on the mutations. It implies that the masking among all positions on the sequence can do accurate predictions of the shared residues but hardly capture the mutation information.

We also conduct AGP and MPP to finetune the baseline model AntiBERT. The pre-training results are shown in Table 8. We can find that without the concatenation of the antibody and its germline, it is difficult to predict the ancestor relationship. It also underperforms than `EATLM` in MPP.

**Table 8:** The pretraining details for different models. The ‘Germline’ is the prediction accuracy of AGP, and the ‘Position’ and ‘Mutation’ are the accuracy of mutation positions and the mutated residues respectively.

| Model | Size | Loss | Step | Germline | Position | Mutation |
| --- | --- | --- | --- | --- | --- | --- |
| AntiBERT | 85M | 0.437 | 568000 | / | / | / |
| AntiBERT w AGP | 85M | 0.558 | 587000 | 0.330 | / | / |
| AntiBERT w MPP | 85M | 1.744 | 622000 | / | 0.965 | 0.410 |
| ESM-1 FT | 85M | 0.2636 | 136000 | / | / | / |
| ESM-1b FT | 650M | 0.2704 | 278000 | / | / | / |
| EATLM w/o AGP & MPP | 85M | 0.143 | 108000 | / | / | / |
| EATLM w/o AGP | 85M | 1.744 | 134000 | / | 1.000 | 0.443 |
| EATLM w/o MPP | 85M | 0.001 | 126000 | 1.000 | / | / |
| EATLM | 85M | 1.856 | 252500 | 0.9960 | 1.000 | 0.443 |
| EATLM-large w/o AGP & MPP | 650M | 0.149 | 193000 | / | / | / |
| EATLM-large w/o AGP | 650M | 1.753 | 267000 | / | 1.000 | 0.438 |
| EATLM-large w/o MPP | 650M | 0.001 | 200000 | 1.000 | / | / |
| EATLM-large | 650M | 1.773 | 384000 | 0.996 | 1.000 | 0.431 |

##### Negative sampling ratio

We have tried the ratio of 0.1/0.3/0.5/0.75 and found that this ratio has little influence on performance and convergence speed. As we discuss in the appendix, the model can quickly converge for AGP and get an accuracy of 0.99.

##### Finetuned Protein Language Models and Larger Architecture

We pre-train our method with a larger architecture and compare it with ESM-1b, which also has 650M parameters. We also further pre-trained the ESMs to transfer to the antibody field. After that, we evaluate them on the antigen binding and paratope prediction task. The results are shown in Table 9. The result show that the larger architecture does show advantage in terms of the performance improvement. For antigen binding, ESM-1b has better performance than ESM-1. However, for paratope prediction, it performs worse. In addition, for ESM, fine-tuning of the antibody dataset may cause the overfitting problem, leading to the decrease in the performance of all three tasks.

**Table 9:** The performance of larger models and finetuned protein language models on three tasks. The ESM-1 and ESM-1b models have 85M and 650M respectively. The `EATLM`-large have similar architecture with ESM-1b. ‘FT’ indicates the model is further finetuned on our antibody pre-training dataset.

|  | Antigen Binding |  |  | Paratope |  |  |
| --- | --- | --- | --- | --- | --- | --- |
|  | AUC | F1 | MCC | AUC | F1 | MCC |
| ESM-1 | 0.917+0.001 | 0.854+0.002 | 0.689+0.002 | 0.886+0.009 | 0.669+0.024 | 0.547+0.026 |
| ESM-1 FT | 0.914+0.001 | 0.858+0.001 | 0.695+0.003 | 0.883+0.009 | 0.657+0.031 | 0.534+0.030 |
| ESM-1b | **0.924+0.002** | 0.860+0.002 | **0.707+0.003** | 0.873+0.009 | 0.655+0.042 | 0.524+0.036 |
| ESM-1b FT | 0.916+0.003 | 0.844+0.003 | 0.686+0.004 | 0.869+0.010 | 0.660+0.021 | 0.527+0.015 |
| EATLM | 0.922+0.052 | **0.862+0.021** | 0.699+0.027 | 0.887+0.008 | **0.698+0.017** | **0.575+0.024** |
| EATLM-large | 0.921+0.004 | 0.854+0.004 | 0.677+0.010 | **0.887+0.007** | 0.685+0.015 | 0.561+0.020 |

### A.6 Discussion about of EATLM

First, EATLM doesn’t use any 3D structure information during pre-training. As a special subgroup of proteins, antibody structures provide much more information such as geometry than sequences. In the future, recruiting structure information for antibody pre-training may be able to improve the results. However, the data scale available for antibody structure is dramatically less than that of antibody sequences. The largest dataset of antibody structures only contains thousands of 3D high-resolution structures, while the number of antibody sequences is in billions. Using structure prediction methods like AlphaFold may help to bridge the gap between sequences and structures. Second, EATLM requires germline as input during downstream tasks, this will slow down the prediction speed.

### A.7 New SARS Binder Discovery

The main challenge for disease diagnosis is to distinguish the disease-related antibodies from millions of antibody sequences in the individual profile, as stated in Section A.3. Here, with the help of a sequence-level predictor, we can give each sequence a most likely label to help antibody discover, whose accuracy has been verified by the excellent results on individual prediction, which may accelerate the discovery of new antibody sequences.

#### SARS Sequence-level Predictor

As described in Section A.3, we first train a sequence-level predictor for SARS-CoV-2. The results are shown in Table 10. Compared with Compared with Figure 4 in the main text, we find that good results in the sequence-level predictor do not necessarily mean good results in the antibody discovery. It can be mainly affected by the noisy label of the sequence level.

**Table 10:** Sequence-level predictor for SARS-CoV-2.

| Sequence-level | SARS |  |  |
| --- | --- | --- | --- |
|  | AUC | F1 | MCC |
| No pretrain | 0.894±0.029 | 0.801±0.045 | 0.637±0.078 |
| ESM-1 | 0.903±0.061 | 0.799±0.032 | 0.648±0.093 |
| MSA-1b | 0.914±0.025 | 0.810±0.050 | **0.671±0.080** |
| Ablang-H | 0.915±0.027 | **0.817±0.038** | 0.668±0.068 |
| Ablang-L | 0.893±0.035 | 0.801±0.054 | 0.637±0.099 |
| AntiBERT | **0.916±0.026** | 0.810±0.033 | 0.661±0.060 |
| EATLM | 0.904±0.027 | 0.808±0.035 | 0.643±0.069 |
| EATLM w/o AGP | 0.904±0.029 | 0.808±0.039 | 0.644±0.078 |
| EATLM w/o MPP | 0.901±0.026 | 0.808±0.035 | 0.648±0.069 |
| EATLM w/o MPP & AGP | 0.901±0.026 | 0.807±0.034 | 0.645±0.067 |

#### Figure out SARS Binders

As shown in Table 3 in the main body, we find 2 true SARS binders and 9 potential binders with the help of `EATLM`. Specifically, we first use our sequence-level predictor to get a probability score for each sequence in the SARS dataset. Then we select the sequence with high-ranked score (the probability > 0.5) and compare them with the public Cov-AbDab database Raybould et al. (2021) ^1^, which contains data on published/patented antibodies known to bind to SARS-CoV-2 (Raybould et al., 2021). Since the CDR3 fragment in the heavy chain is the most relevant to the binding of antibody and antigen, we calculate the edit distance between the CDR3 fragments in heavy chains (CDR-H3) with those of the known binder and use a threshold of 85% similarity as the sequence identity. 85% Hamming distance for B cell antibody sequence clustering (identify similar B cell antibody sequences responding to the same antigen/epitope) was previously suggested in this paper (Gupta et al., 2017). This method then was widely used for B cell antibody repertoire analysis in different studies (Montague et al., 2021; Wang et al., 2022).

#### SARS Binder Analysis

To provide a more intuitive analysis of the similarity between our predicted antibody and true SARS-CoV-2 binders, we investigate the 3D structure of the true binding antibodies and the mutation site of our predicted sequence on the corresponding structure. High resolution structure of true binding antibody #3 in Table 3 with SARS-CoV-2 are shown in Figure 13 (PDB code: 7N62). The interaction interface between the antibodies and SARS-CoV-2 spike/RBD are shown in Figure 3 in main body. CDR-H3 were shown in orange. Only one single atom highlighted in red is different between predicted binder and true binder. Obviously, these different residues don’t localize to direct binding site and CDR-H3 founding core, suggesting the sequence difference likely will not affect antibody virus interaction. Furthermore, we found the epitopes of the 11 identified SARS-CoV-2 antibodies cover a wide range of different structures from traditional RBD domain to novel non-RBD epitopes like S2 and NTD as shown in Table 3. This result shows our method enables diverse-epitope antibody discovery.

#### Probability Threshold Sensitivity

In order to investigate the influence of the threshold used to determine the potential binders, we try different thresholds in Table 11. Here, the probability threshold means that if the sequence predictor give a probability higher than the threshold for one sequence, it will be viewed as a potential binder. If the predicted binder have a sequence similarity higher than 85% with the existing binders in Cov-AbDab, we view it as one hit. As the threshold score increases, the hit rate corresponding increases from 0.528% to 0.562%, indicating that our model may enable priority selection of SARS-CoV-2 antibodies and reduce experimental costs.

**Table 11:** SARS-CoV-2 antibody hit rate with different probability thresholds. ‘Total’ is the total number of sequences whose probabilities are higher than the threshold. ‘Hit’ is the number of sequences that meet the similarity requirements with existing binders.

| Threshold | Total | Hit | Hit rate (%) |
| --- | --- | --- | --- |
| 0.5 | 13253 | 66 | 0.498 |
| 0.7 | 10227 | 54 | 0.528 |
| 0.8 | 9338 | 49 | 0.525 |
| 0.9 | 8178 | 47 | 0.562 |

**Table 12:**
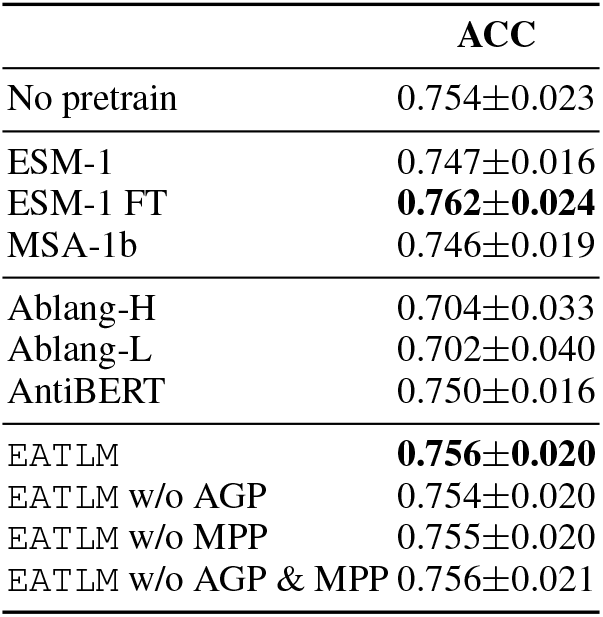
The Q7 disease classification task. ‘ACC’ is the accuracy rate.

|  | ACC |
| --- | --- |
| No pretrain | 0.754±0.023 |
| ESM-1 | 0.747±0.016 |
| ESM-1 FT | **0.762±0.024** |
| MSA-1b | 0.746±0.019 |
| Ablang-H | 0.704±0.033 |
| Ablang-L | 0.702±0.040 |
| AntiBERT | 0.750±0.016 |
| EATLM | **0.756±0.020** |
| EATLM w/o AGP | 0.754±0.020 |
| EATLM w/o MPP | 0.755±0.020 |
| EATLM w/o AGP & MPP | 0.756±0.021 |

**Figure 13:**
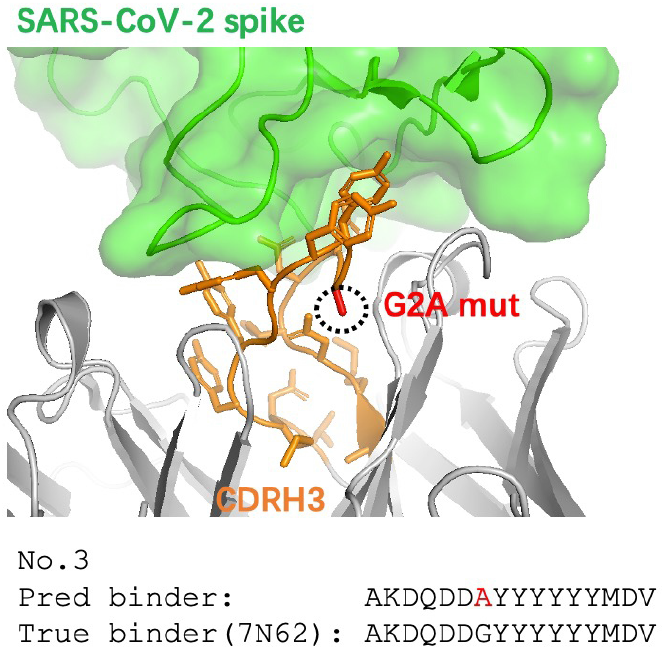
High resolution structure of mutation in the predicted binder (AKDQDDAYYYYYYMDV) with the existing binding antibody (AKDQDDGYYYYYYMDV).

#### Sequence Similarity Sensitivity

In previous work, two antibodies with the CDR-H3 similarity over 85% can be viewed as similar and have a high probability to share the same functionality. And here we also check the influence on the binder matching of different thresholds of the similarity. The results are shown in 14. Here, we fix the probability threshold as 0.5. As we can see, the baselines have similar trends in all threshold. If we relax the threshold, there will be more matching sequences. However, the predictors will have less advantage over the random order, which indicates that the ranking is less important.

#### The Potential of New Binder Discovery

During the training of our sequence-level predictor, we have no reliable ground-truth labels, which means that the model has never known which sequences can bind to SARS in the real-world scenario. However, the model can learn from the noisy data and rank the real SARS binders with high probabilities. Sequence identity of 1 means that the CDR-H3 fragment can be directly found in the Cov-AbDab database, which implies that the sequences have been verified by wet laboratory testing. The other sequences with an identity over 90% are thought to have a similar binding performance to existing binders, indicating that they are promising SARS binders that can help the discovery of therapeutic antibodies for SARS-CoV-2.

**Figure 14:**
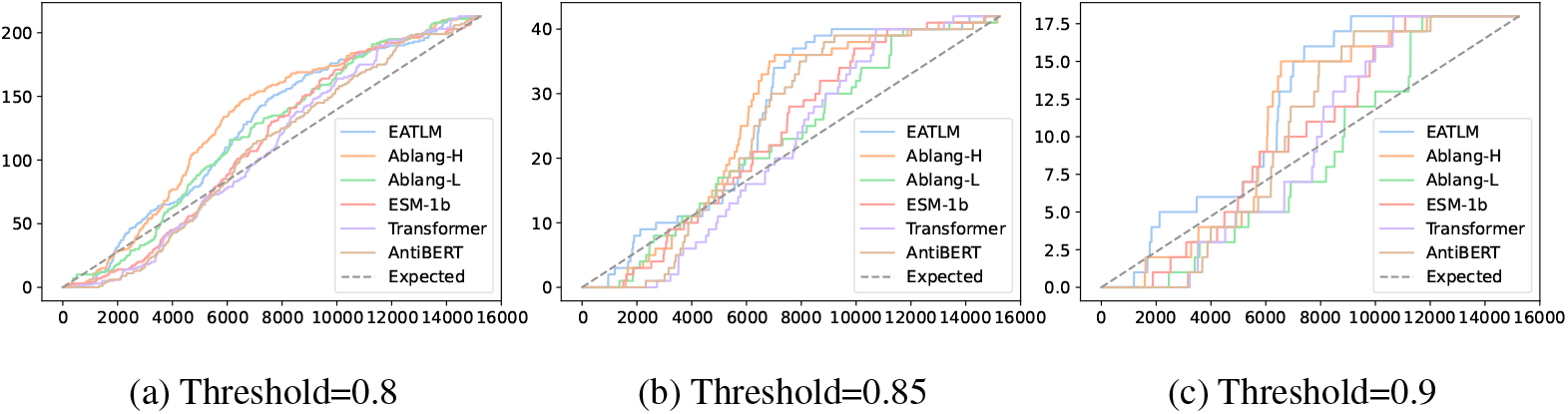
The cumulative count of matching sequences number. The dashed line is the expected results for a random order. The x-axis is the sequence number and the y-axis is the cumulative matched sequence number.

### A.8 Extented Study for Disease Diagnosis

It would be interesting to see whether our sequence classifier can be used for other applications, such as disease diagnosis. Each human is estimated to maintain about **10**^8^ - **10**^10^ distinct antibody sequences, constructing an informative encyclopedia recording the past and present health and disease. Interpreting the pattern of the sequences has already proved useful in disease diagnosis and allows us to assess many infectious diseases without expensive laboratory testing. However, it is difficult to distinguish which antibody sequence from the numerous sequences is responsible for the recognition of the specific antigen, which hinders the discovery of the antibody for diseases (Zaslavsky et al., 2022; Lu et al., 2018; Greiff et al., 2020).

Benefiting from the recent high-throughput sequencing, we can obtain millions of antibody sequences from the individual human. At the same time, we can get a disease label that indicates whether the human is infected by the disease. The main challenge is that we can hardly get the sequence-level disease label that indicates whether the antibody sequence is related to the disease. Thus, we follow the practice of Roskin et al. (2020) to use the individual label as the rough sequence label and train a sequence-level predictor. Then we use this predictor to predict sequences of the individual profile and make the trimmed mean score as the individual score.

We use the same data processing as Antibody Discovery stated in Section A.3. For health/SARS/HIV/Ebola/Allergy/SLE/MS, we set the ‘*Disease*’ field to ‘*None*’, ‘*Ebola*’, ‘*Allgery*’, ‘*SLE*’, ‘*MS*’. Then we obtain 87/133/51/14/12/8/8 patient profiles for each type. We also do 10-cross validation and select sequences with high redundancy.

#### Disease Classification

We use all these diseases profile to build the Q7 classification task for disease diagnosis. Previous biological studies mainly use this multi-classification task for disease diagnosis Zaslavsky et al. (2022); Wang et al. (2022), highlighting the discriminatory power among different diseases are important for disease diagnosis. The results are shown in Table 12. We found both PPLM an PALM show comparable results as the random initialized model, suggesting the finetuning part play more important role and pretrained language model cannot help this task.

#### Sequence-level Predictor for Various Disease

As before, we train a sequence-level predictor for each disease. The results are shown in Table 13. Compared with Table 4 in the main text, we find that good results in the sequence-level predictor do not necessarily mean good results in the individual-level predictor. It is mainly due to the trimmed mean we use to get individual-level results, which is a central estimate that is robust to noise labels. Overall, our model has comparable results to other models in terms of sequence prediction with noisy labels, and has better results for individual diagnosis.

**Table 13:**
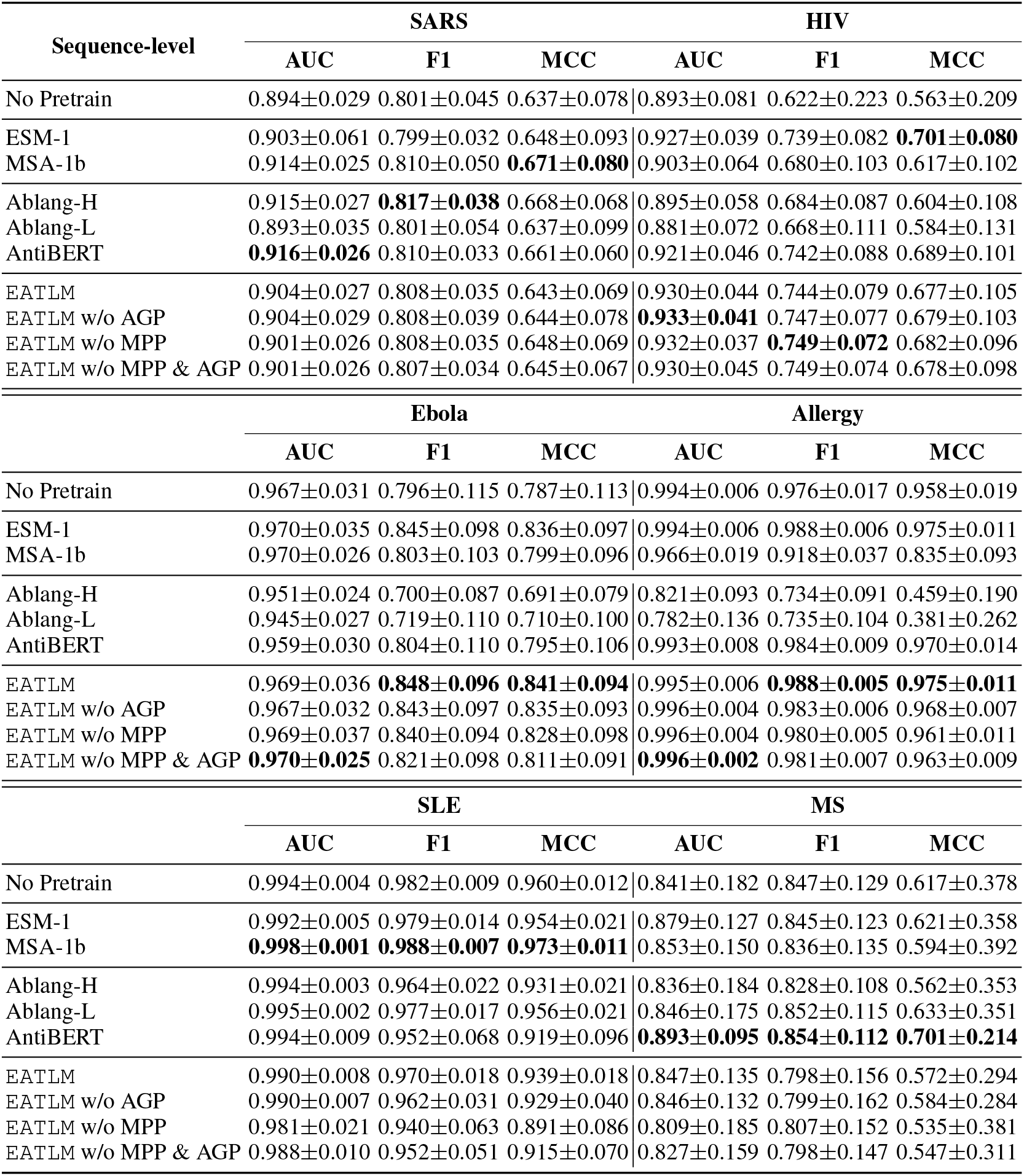
Sequence-level predictor for disease diagnosis.

#### Individual-level Predictor for Various Disease

It is observed our evolution-aware EATLM performs the best in the individual-level classifier to determine whether the patient suffering from SARS. Besides, PALMs significantly outperform PPLMs. The results is shown in Table 14.

**Table 14:** Individual-level predictor for disease diagnosis.

| Pateint-level | SARS |  |  | HIV |  |  |
| --- | --- | --- | --- | --- | --- | --- |
|  | AUC | F1 | MCC | AUC | F1 | MCC |
| No pretrain | 0.975±0.033 | 0.962±0.024 | 0.902±0.062 | 0.920±0.092 | 0.815±0.117 | 0.776±0.122 |
| ESM-1 | 0.979±0.017 | 0.933±0.032 | 0.824±0.090 | 0.967±0.056 | 0.883±0.107 | 0.849±0.135 |
| MSA-1b | 0.989±0.012 | 0.954±0.033 | 0.887±0.078 | 0.962±0.039 | 0.833±0.068 | 0.789±0.072 |
| Ablang-H | 0.988±0.013 | 0.968±0.024 | 0.920±0.063 | 0.960±0.040 | 0.788±0.134 | 0.744±0.137 |
| Ablang-L | **0.990±0.015** | 0.938±0.039 | 0.843±0.091 | 0.931±0.069 | 0.798±0.131 | 0.742±0.170 |
| AntiBERT | 0.983±0.017 | 0.976±0.022 | 0.938±0.056 | 0.969±0.053 | 0.878±0.112 | 0.850±0.134 |
| EATLM | 0.977±0.028 | **0.983±0.017** | **0.955±0.045** | **0.978±0.054** | 0.914±0.081 | 0.884±0.106 |
| EATLM w/o AGP | 0.983±0.019 | 0.973±0.025 | 0.929±0.066 | **0.978±0.054** | 0.903±0.076 | 0.869±0.099 |
| EATLM w/o MPP | 0.987±0.015 | 0.969±0.024 | 0.921±0.061 | 0.976±0.053 | 0.906±0.092 | 0.880±0.112 |
| EATLM w/o MPP & AGP | 0.986±0.014 | 0.969±0.023 | 0.920±0.061 | 0.973±0.052 | **0.919±0.096** | **0.896±0.117** |
|  | Ebola |  |  | Allergy |  |  |
|  | AUC | F1 | MCC | AUC | F1 | MCC |
| No Pretrain | 0.994±0.017 | 0.933±0.133 | 0.934±0.132 | 1.000±0.000 | 1.000±0.000 | 1.000±0.000 |
| ESM-1 | 1.000±0.000 | 0.933±0.133 | 0.934±0.132 | 1.000±0.000 | 1.000±0.000 | 1.000±0.000 |
| MSA-1b | 1.000±0.000 | 0.933±0.133 | 0.934±0.132 | 1.000±0.000 | 1.000±0.000 | 1.000±0.000 |
| Ablang-H | 0.944±0.167 | 0.767±0.213 | 0.750±0.254 | 1.000±0.000 | 0.960±0.080 | 0.916±0.169 |
| Ablang-L | 0.961±0.100 | 0.731±0.205 | 0.703±0.277 | 1.000±0.000 | 0.960±0.080 | 0.916±0.169 |
| AntiBERT | 0.978±0.067 | 0.933±0.133 | 0.934±0.132 | 1.000±0.000 | 1.000±0.000 | 1.000±0.000 |
| EATLM | 1.000±0.000 | **0.947±0.111** | **0.944±0.114** | 1.000±0.000 | 1.000±0.000 | 1.000±0.000 |
| EATLM w/o AGP | 1.000±0.000 | 0.933±0.133 | 0.934±0.132 | 1.000±0.000 | 1.000±0.000 | 1.000±0.000 |
| EATLM w/o MPP | 1.000±0.000 | 0.933±0.133 | 0.934±0.132 | 1.000±0.000 | 1.000±0.000 | 1.000±0.000 |
| EATLM w/o MPP & AGP | 0.994±0.017 | 0.933±0.133 | 0.934±0.132 | 1.000±0.000 | 1.000±0.000 | 1.000±0.000 |
|  | SLE |  |  | MS |  |  |
|  | AUC | F1 | MCC | AUC | F1 | MCC |
| No Pretrain | 1.000±0.000 | 1.000±0.000 | 1.000±0.000 | 0.700±0.200 | 0.900±0.137 | 0.893±0.400 |
| ESM-1 | 1.000±0.000 | 1.000±0.000 | 1.000±0.000 | 0.900±0.200 | 0.933±0.133 | 0.900±0.200 |
| MSA-1b | 1.000±0.000 | 1.000±0.000 | 1.000±0.000 | 0.700±0.400 | 0.893±0.137 | 0.700±0.400 |
| Ablang-H | 1.000±0.000 | 1.000±0.000 | 1.000±0.000 | 0.800±0.245 | 0.893±0.137 | 0.700±0.400 |
| Ablang-L | 1.000±0.000 | 1.000±0.000 | 1.000±0.000 | 1.000±0.000 | 0.960±0.080 | 0.800±0.400 |
| AntiBERT | 1.000±0.000 | 1.000±0.000 | 1.000±0.000 | 1.000±0.000 | 1.000±0.000 | 1.000±0.000 |
| EATLM | 1.000±0.000 | 1.000±0.000 | 1.000±0.000 | 1.000±0.000 | 0.867±0.163 | 0.800±0.245 |
| EATLM w/o AGP | 1.000±0.000 | 1.000±0.000 | 1.000±0.000 | 1.000±0.000 | 0.867±0.163 | 0.800±0.245 |
| EATLM w/o MPP | 1.000±0.000 | 1.000±0.000 | 1.000±0.000 | 1.000±0.000 | 0.827±0.150 | 0.600±0.374 |
| EATLM w/o MPP & AGP | 1.000±0.000 | 1.000±0.000 | 1.000±0.000 | 1.000±0.000 | 0.827±0.150 | 0.800±0.374 |

## Footnotes

1 http://opig.stats.ox.ac.uk/webapps/covabdab/

## Notes

https://github.com/dqwang122/EATLM

### Competing Interest Statement

The authors have declared no competing interest.

